# Climate Resilience Index as a tool to explore households’ resilience to climate change-induced shocks in Dinki watershed, central highlands of Ethiopia Households’ resilience to climate change impacts

**DOI:** 10.1101/382358

**Authors:** Mengistu Asmamaw, Argaw Ambelu, Seid Tiku Mereta

## Abstract

This study assessed households’ resilience to climate change-induced shocks in Dinki watershed, northcentral highlands of Ethiopia. The data were collected through cross-sectional survey conducted on 288 households, six focus group discussions and 15 key informant interviews. The Climate Resilience Index (CRI) and the Livelihood Resilience Index (LRI) based on the three-resilience capacities (3Ds) frame, using absorptive, adaptive and transformative, were used to measure households’ resilience to climate change-induced shocks on agro-ecological unit of analysis. Findings indicate that the CRI and the resilience capacities based on the indexed scores of major components clearly differentiated the study communities in terms of their agro-ecological zones. Specifically, the LRI score showed that absorptive capacity (0.495) was the leading contributing factor to resilience followed by adaptive (0.449) and transformative (0.387) capacities. Likewise, the midland was relatively more resilient with a mean index value of 0.461. The study showed that access to and use of livelihood resources, such as farmlands and livestock holdings, diversity of income sources, infrastructure and social capital were determinants of households’ resilience. In general, it might be due to their exposure to recurrent shocks coupled with limited adaptive capacities including underdeveloped public services, poor livelihood diversification practices, among others, the study communities showed minimal resilience capacity with a mean score of 0.44. Thus, in addition to short-term buffering strategies, intervention priority focusing on both adaptive and transformative capacities, particularly focusing on most vulnerable localities and constrained livelihood strategies, would contribute to ensure long-term resilience in the study communities.

## 1. Introduction

Climate change-induced shocks are the major livelihood threats of humanity, where underdeveloped countries are disproportionately hit by adverse effects (1). The projections by the Intergovernmental Panel on Climate Change (IPCC) shows that the frequency and intensity of climate change-induced shocks, such as heat waves, droughts, floods, etc. are growing all over the world (2). The effects of such extreme weather events would add extra stress on human health, food security and water resources, where the rural poor are extremely susceptible and adversely impacted [3, 2]. The IPCC report emphasized that disaster risk management programs should focus on reducing exposure and vulnerability while enhancing resilience to shock impacts (2).

The concept of resilience stems to the Latin ‘resilire’ to denote to ‘bouncing back’ or ‘recoiling’ (4). The term was primarily applied in mechanics in 1858 to denote the capability of a material to resist a force (rigidity) as well as to absorb the force with deformation; later it was used in psychology in 1950s, in system ecology in 1973 and in social-ecological systems in 1990s (4). The intensification of two huge societal trends-climate change and globalization, which amplify multifaceted and non-directional impacts have caused resilience to be acknowledged in wide range of disciplines globally (5). Aiming to address the overwhelming environmental issues, such as disaster risk reduction, climate change adaptation, vulnerability, social protection, etc.(6), its application has gradually expanded into social-ecological system and defined as the potential of a social-ecological system to sustain basic structures and continue functioning following shock events [7, 8]. Being a multidisciplinary term, it has been applied in diversity of connotations, yet all share common point on ‘the ability to respond to changes, particularly unprecedented changes’(5).

The capability of a social-ecological system to respond to extreme shock events encompasses multiplicity of abilities including “shock absorbing”, “buffering”, “bouncing back”, and “transforming” (8). Its application in various disciples has broaden its understanding from its original narrowed engineering resilience- ‘the potential of a system to bounce back after disturbance’ into more comprehensive concept- ‘the ability not only to bounce back but also to adapt to and even to transform into new system’ (6). Furthermore, a socio-ecological resilience is perceived as a process than a static state and should acquire and maintain the three-core resilience capacities, namely absorptive, adaptive and transformative to sustain long-term resilience [6, 10]. As absorptive, adaptive and transformative capacities are considered as the three major structural elements and best to capture resilience (6), this study followed the three-capacities (absorptive, adaptive and transformative capacities) frame to explore households’ resilience to climate change-induced shocks.

The three core responses or resilience capacities can be linked depending on shock intensity. Accordingly, during minimal shock incidence, it is natural that the system would block or resist it (6). Hence internal resistance is known as the natural characteristic of a system manifested on daily basis where resources could block the shock enabling the system to continue functioning-highly comparable to the human immune system (9). Absorptive capacity is especially basis to buffer short-term disturbances as well as during the beginning phase of coping of huge shocks (5).

The next adaptive resilience involving system adjustment to sustain system functioning will be exercised if the shock exceeded the absorptive capacity (10). Adaptive capacity is “the ability of a system to adjust itself to sustain system functioning”(11). These adjustment practices are incremental as well as learning through failure and success that adds to adaptability (12). This capacity involves “resourcefulness-the potential to identify challenges, develop priorities, mobilize resources, to integrate experience and knowledge during crises, to plan for upcoming shock impacts” (5). These multi-level (individuals, households, community) and incremental adjustment mechanisms for farming communities may include livelihood diversification, establishing market networks, empowering storage facilities, developing pooling among communities, introducing of shock resistance varieties, new farming practices, strengthening social networks, etc. (6).

In the case of high intensity and recurrent shocks, it may be difficult to sustain system functioning through adaptive resilience, involving transformative resilience. It is often associated to system-level changes in factors like infrastructure (example: road, communication, credit access, health facilities, etc.), governance, formal safety nets which substantially strengthen long-term resilience (13). For instance, changing of the agrarian livelihood into resource extraction economy, ecotourism, change in resource management practices, etc. Transformative response may require institutional reforms, behavioral changes and technological innovations (14). Factors like socioeconomic policies, land-use policies, resource management trends, institutions and technology may limit the performance of transformative resilience (14).

In the face of environmental uncertainty, households’ capacities to effectively respond to the alarmingly growing shock events needs to be strengthened (5) to enable smallholder farmers to better withstand the upcoming shock impacts (15). Because resilient households are more active to anticipate, resist, cope with and recover against shock impacts (16) as well as to sustain or improve standard of living in the face of environemntal changes (17). The findings of the study would help to prioritize intervention measures for livelihood resilience by identifying adaptation limits in Dinki watershed, northcentral highlands of Ethiopia.

## 2. Materials and Methods

### 2.1 The Study Area

Dinki watershed is found in Ankober district in central highlands of Ethiopia (Fig. 1). Ankober is located between 9° 22’-9° 45’ N latitude and 039° 40’-039° 53’ E longitude (Fig. 1). Most of the district area are hills and mountainous (75%), where rugged terrains and plain topography account for 17% and 8%, respectively. More than half of the district (53%) has *woinadega* (equivalent to sub-tropical), climatic condition followed by *kola* (tropical) climate; where dega (temperate) and wurich (cool) climates constitute 10.5 and 1.5 percent, respectively (18). The Rainfall pattern is bimodal where some short and long-term rainy periods are recorded in March and in late June to September, respectively. A 30 year (1987-2016) of metrological data showed a mean annual rainfall of 1,179 mm; where the mean minimum and maximum monthly temperature was 6.47 and 19.99 °C, respectively.

**Figure 1.**
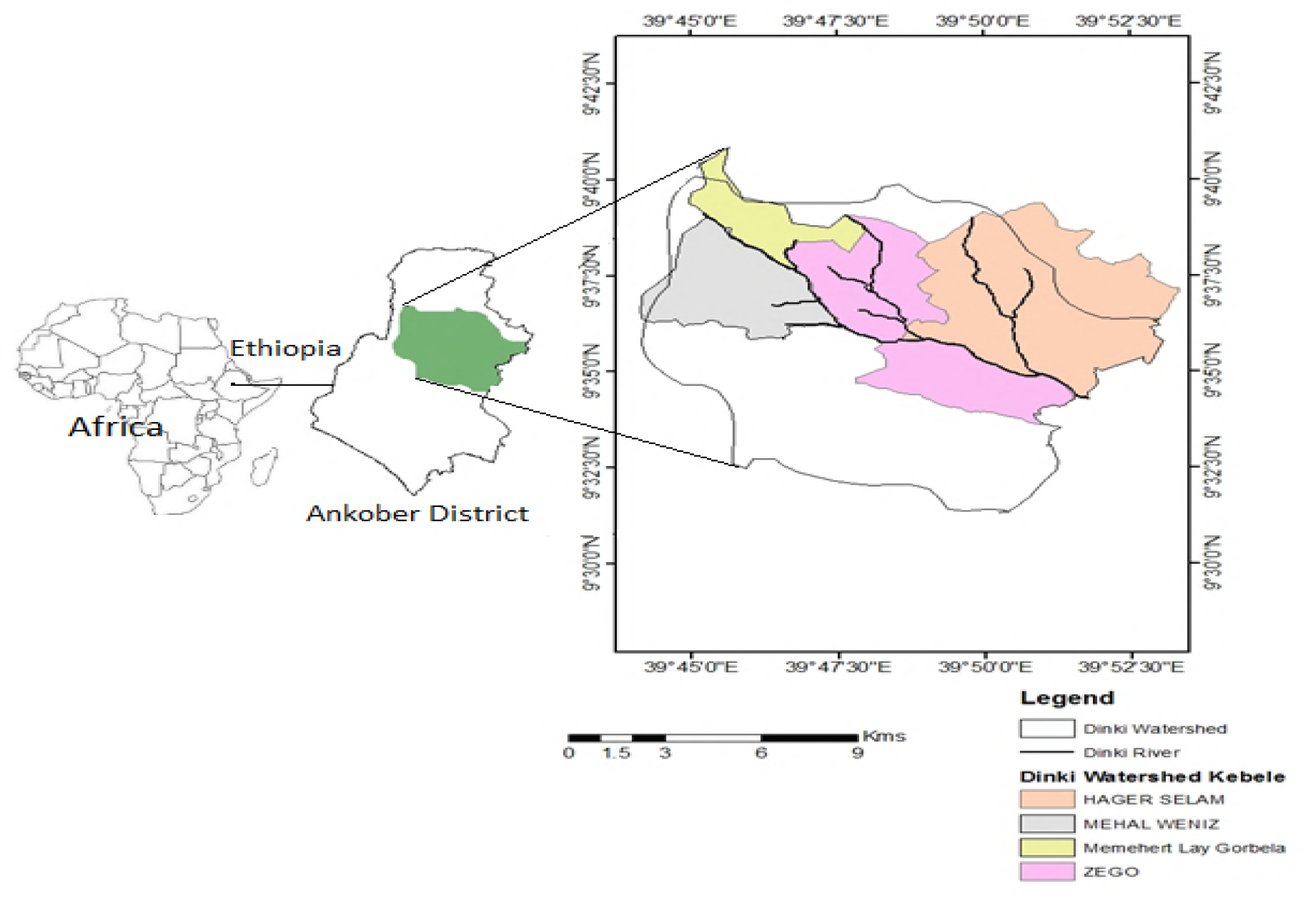
Map of the Dinki watershed, central highlands of Ethiopia

### 2.2 Data Collection Techniques

Data were collected through participatory rural appraisal between December 2017 to February 2018 through focus group discussion, key informant interview and household survey viz: highland, midland and lowland agro-climatic zones (AEZs). Prior to data collection, an ethical clearance letter was received from the Institutional Review Board (IRB), institute of Health, Jimma university.

#### 2.2.1 Qualitative data collection

Six focus group discussions (FGDs) (two gender-segregated FGDs in each agro-ecology), each comprising 8-12 participants were conducted to collect data on livelihood vulnerability and resilience to climate change-induced shocks. Some of the questions asked were: What socioeconomic and environmental factors do you think determine resilience in this locality? Do you think inter-household resilience variability in this locality? What experience are there in this locality to prepare for, mitigate with and cope with (absorptive capacities); adjust to sustain system functioning (adaptive capacities) and strengthen long-term resilience like through system changes in land-use, natural resource management, governance, etc. (transformative capacities)?

The same interview questions were used to conduct 15 face-to-face interviews involving various community members, such as religious leaders, watershed management group members, elders, youth, women as well as representatives from school and development agents to explore the incidences of climate change-induced shocks and adaptation strategies contributing to manage disturbances. A snowball approach was used to purposively select participants for interview and information redundancy was used as an insurance for information saturation.

#### 2.2.2 Quantitative data collection

Based on the feedback and information from qualitative data, a standardized questionnaire was developed. In addition to the questions used in the interviews, a sample of questions asked in the questionnaire survey were: What do you think the resilience status of this locality? Was there any environmental and/or socioeconomic shock during the last 12 months? Do you think climate change-induced shocks affect your livelihood strategies? What coping strategies do you use to prepare for, mitigate with or prevent the negative impacts of shocks? What adjustment strategies (example: livelihood diversification, farming practice, social networking, etc.) do you apply to sustain system functioning even during crises? Is there any system-level change (example: infrastructure, governance, social networking, etc.) that supports to strengthen long-term resilience in this locality? A simple random sampling technique was employed to select 294 respondents from a total of 1,245 households; where prescriptions by Kothari (19)was used to calculate the sample size.

### 2.3 Climate Resilience Index (CRI) calculation

As resilience is a complex concept, its quantification remains debatable. Currently, however, proxy indicators through composite index frame has been used to measure resilience in wide range of literature [21, 22]. The climate resilience index (CRI) development followed the prescription by Tambo (21). Accordingly, a tool developed by FAO (20) to measure food insecurity was customized to assess households’ resilience to climate change-induced shocks. The tool consists of ten major components and a household with higher in average values of each component is hypothesized to be resilient to climate change-induced shocks. Stakeholders consultation (extension workers, development agents, experts and elders) and review literature [21, 10, 22] were used to select relevant indicator and the details are presented in Table 1 below.

**Table 1.**
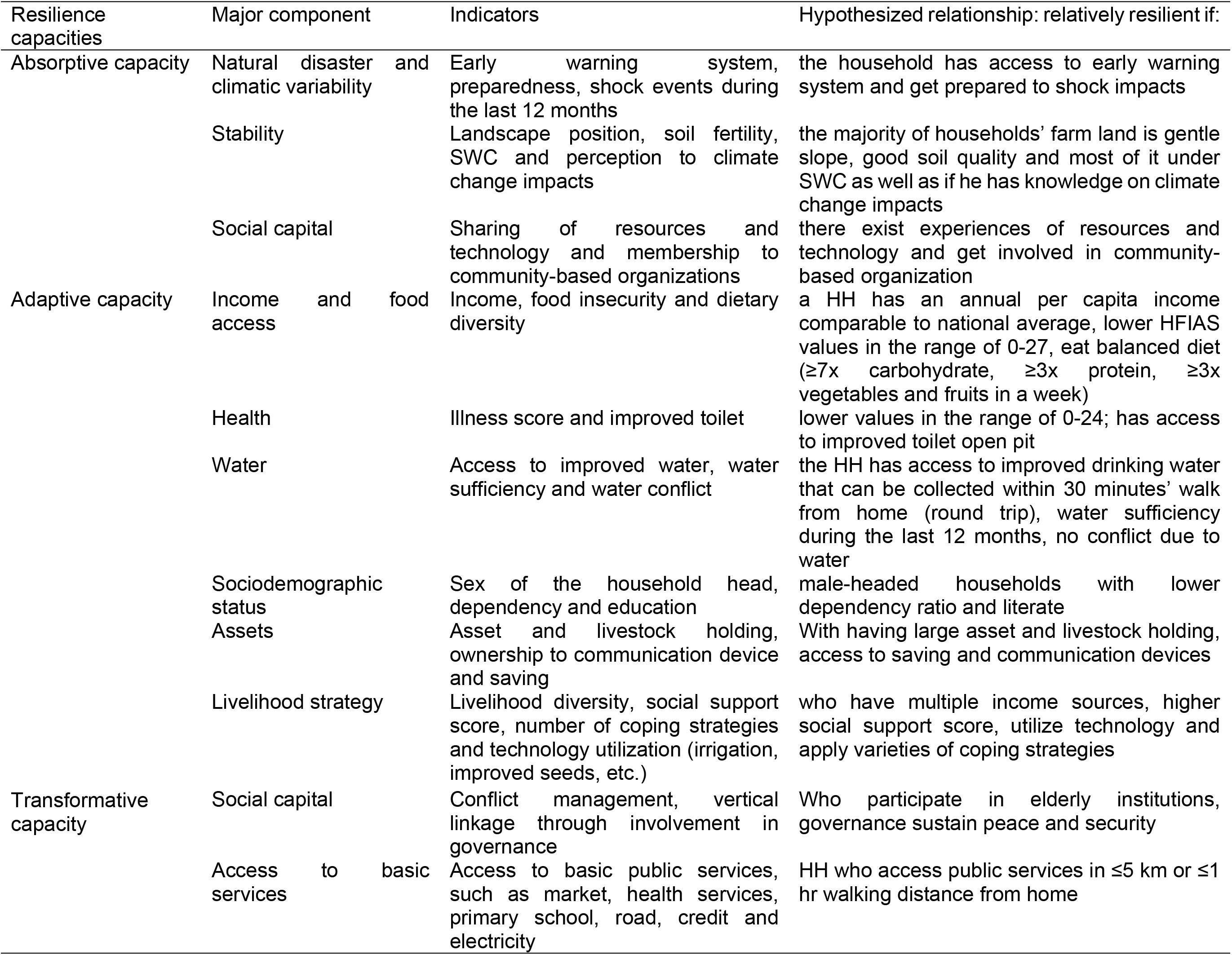
Resilience capacities, major components, sub-components and hypothesized relationships

The CRI uses a balanced weighted technique (23) where each sub-component (indicator) contributes equally to the index. Using a household-level data on these indicators, a Climate Resilience Index (CRI) was developed on agro-ecological unit of analysis. As each major component is composed of different number of indicators measured on different scales, the standardization considered the functional relationship between indicators and resilience (21). In effect, two methods of standardization were employed. Indicators that are expected to have direct relationship with resilience, such as income and food access, diversity of income sources, coping strategies, etc. were standardized using equation (1) as:

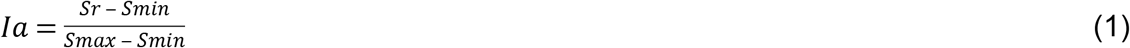

Whereas indicators expected to have inversely related to resilience, such as household food insecurity and access score (HFIAs), illness score, shock events, etc. were standardized using equation (2):

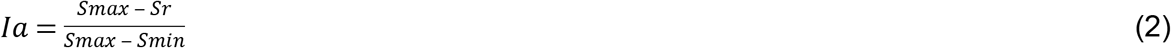

Where Ia is the standardized value for the indicator a, Sr is the observed (average) value of the indicator for agro-ecology r, min and max are the minimum and maximum values of the indicator across all the agro-ecology, respectively. Once each indicator has been standardized, the average value of each major component was computed using equation 3:

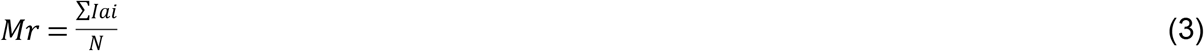

Where Mr is one of the ten major components for agro-ecology r, Iai is the indicator indexed by i, that make up each major component, N is the number of indicator in each major component. After values for each of the ten major components for each agro-ecology were calculated, the CRI was obtained from the weighted average of the ten components as:

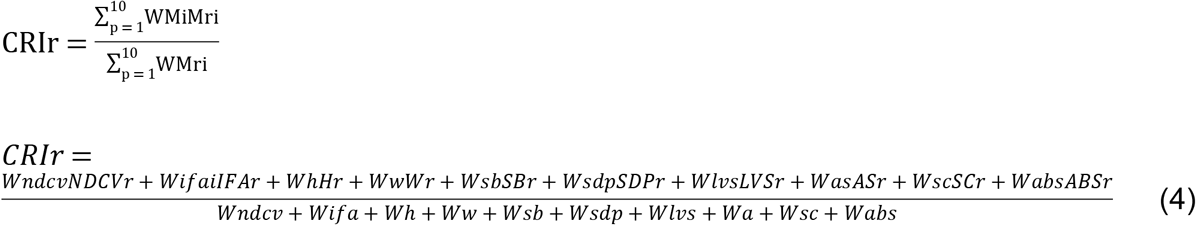

Where CRIr is the Climate Resilience Index for each agro-ecological zone, Mri= the number of indicators of the major component, WMi= weight of major component i, NDCV=natural disaster and climate variability, IFA= income and food access, H=health, W=water, Sb=stability, SDP=sociodemographic profile, LVS=livelihood strategy, A=assets, SC=social capital, ABS=access to basic services.

In order to better understand resilience, the Climate Resilience Index (CRI) frame indicators were aggregated into the three resilience capacities (3Ds) viz: absorptive, adaptive and transformative capacities [6, 10, 5, 25] absorptive capacity is the ability of a socio-ecological system to prepare for, mitigate with or prevent negative impacts through coping strategies in order to preserve and restore basic structures and functions (24). The index was computed based on the perceived ability of households to climate change-induced shocks, access to early warning system, preparedness, stability and social capital like sharing of resources, technology and membership to community-based organizations (13).

Adaptive capacity is the ability of a system to adjust impacts to moderate potential damage, to take advantage of opportunity, so that it continues functioning without significant change in system structures (3). Examples include, livelihood diversification, introducing drought resistant seed varieties (like growing of *Vigna radiate* or mung bean/ *masho*). In effect, income and food access, assets, livelihood diversification strategies, etc. were placed under adaptive capacity [10, 26]. Transformative capacity is the ability to create an enabling new system in times of crises (7). It is often associated to system-level changes in factors like infrastructure (example: road, communication, credit access, health facilities, etc.), governance, formal safety nets which substantially strengthen long-term resilience. As a result, access to basic services, social capital like conflict management mechanisms and vertical linkages were captured under transformative capacity [10, 26]. Therefore, indicators presented in equation (4) were aggregated into respective resilience capacities to generate the livelihood resilience index (LRI) as follows:

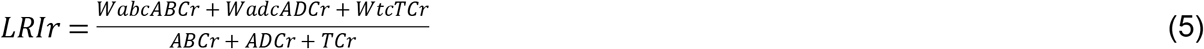

Where LRIr is the resilience index for the agro-ecology r; Wabc, wadc and wtc are the weight of absorptive, adaptive and transformative capacities, respectively; ABCr, ADCr and TCr are the number of indicators in absorptive, adaptive and transformative capacities in each agro-ecological zone, respectively.

## 3. Results and Discussion

### 3.1 Resilience indicators identified in Dinki watershed socio-ecological system

The study households perceive resilience as a state of recovery against climate change-induced shocks without significant help from external institutions. The effects of climate change-induced shocks and consecutive rate of recovery are not uniform across households. In effect, households in Dinki watershed socio-ecological system were classified into poor resilient, moderately resilient and resilient based on their recovery time to climate change-induced shocks. Such classification was also reported in other parts of Ethiopia (25). Key determinants of resilience and major features of each resilience category are presented in Table 2 below.

**Table 2.**
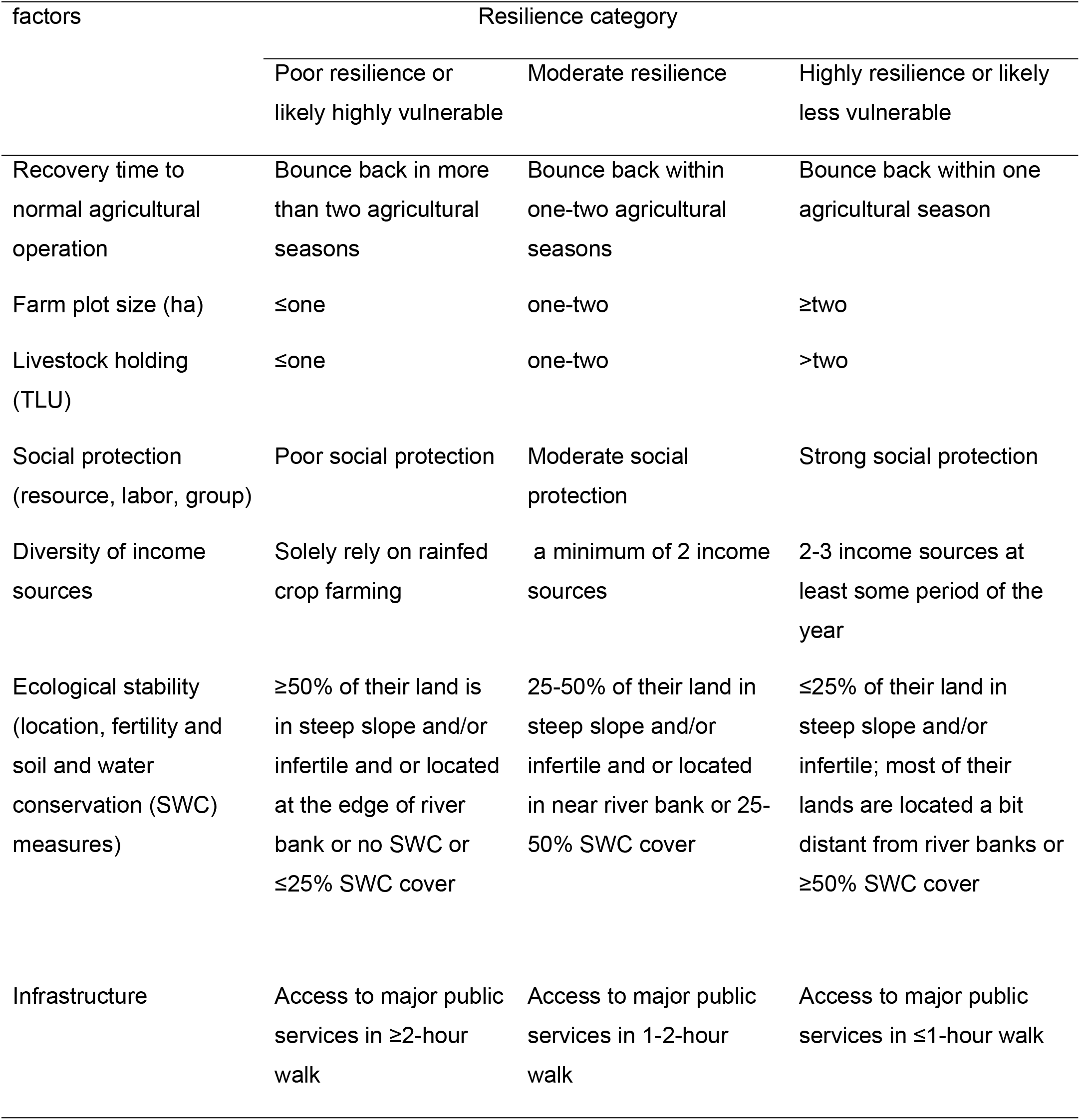
Resilience categories and factors influencing households’ resilience to climate change-induced shocks in Dinki watershed socio-ecological system.

Discussant noted that access to and size of farmland to be determinants of household’ livelihood and resilience to shock impacts. They stressed that land ownership is a priority for farming community for long-term decision and soil fertility management options. Accordingly, landless households are less likely to work on natural resource management practices even may amplify environmental degradation through overexploitation. Whereas, households with large farm sizes are more likely to invest on land and soil fertility management works, diversify income sources (crop-livestock integration, polyculture, agroforestry, etc.) and more likely to bounce back quickly against shock impacts. In agreement with this finding, studies state that landlessness and small land holding are determinant factors causing land degradation and resilience erosion (25). Besides, a study in central Ethiopia discloses that natural resource management practices, which in turn determined by farm size, among others, are strategies for rural communities to enhance their resilience to shock (26).

Livestock holding is argued to signify wealth and dignity in rural Ethiopia. Discussants disclosed that livestock ownership is a determining factor for household livelihood and sustainability; as households having domestic animals are more likely to enhance and diversify income sources. However, the number and diversity of animals critically influence their economic returns. Accordingly, Oxen ownership is a priority for every farmer to secure his agricultural production. The next priority is reported to have milking cow to sustain livestock production and dietary diversity. Depending on the agro-ecology and households’ choice, having of transportation animals, such as donkey/horse/mule/camel would be the next interest. Because, in areas with limited car access, like the study area, humans and materials, including agricultural inputs (fertilizer, improved seeds, pesticides), market inputs and related commodities are transported through these animals; markedly supporting livelihood options, asset accumulation and recovery to shocks. As a result, households with more than two TLU (a minimum of 2 Oxen or 1 Ox + 1 cow) are more likely to be resilient to climate change-induced shocks. In line with this finding, a study in other parts of Ethiopia states that asset holding, including land and livestock unit, is determinant to diversify income sources, improves income and critical for the households’ resilience to food insecurity (27).

Participants disclose that social networking is a determinant factor for mankind to share labor and resources, manage disputes as well as to mitigate with, adapt to and quickly recover against shock impacts. See also [10, 29, 26, 30]. In terms of ecological stability, discussants disclosed that households whose farm lands are located in steep slopes and near to river banks are highly vulnerable to soil erosion and flooding impacts. Likewise, land fertility is also reported as a principal factor influencing households’ productivity and wealth status. Accordingly, households whose farm lands are in gentle slope and with better soil fertility are better off in production and are relatively resilient to shock impacts than their counter parts.

Moreover, soil and water conservation practices are identified as determinant factors affecting households’ resilience to erosion. In effect, households who experience intensive soil and water conservation measures are less likely impacted by erosion and more likely to recover quickly against the adverse impacts of erosion. Thus, poor households are those whose most of their lands are located in steep slopes, proximate to river banks, with infertile and minimal soil and water conservation practices and thereby less resilient to shock impacts. In line with this finding, studies disclose that land location and fertility are critical to determine farm productivity. Accordingly, households with improved land fertility are better off in farm production and more resilient to shocks [27, 26].

#### Diversity of income sources

Discussants and key informants disclosed that households who experience multiple livelihood options have more assets and improved living standards. In this aspect, female discussants stated that small-scale irrigation, home garden and small-scale trading are essential in supporting the income-generating ability of women and youth. Two female informants in *Mehal-Wonz* and *Zego* sites disclosed that selling of alcohol, locally termed as *tela* and *areki* has substantial contribution in improving their standard of living, especially in fulfilling children’s demands of clothing and stationary materials. In general, households with diversity of income sources are less vulnerable; instead more likely quickly recover against climate change-induced shocks than who solely depend on single source of income. In agreement with this finding, studies state that income diversification is a strategy to improve income-generating ability of women in rural households (28). As a result, livelihood diversification is attributed with both coping strategy to risks in times of hazard events, as well as a means of livelihood development in conducive economic settings (29).

#### Access to basic infrastructure

key informants noted that infrastructure, mainly road and market are basis for further societal developments. In this aspect, access to basic infrastructures is minimal where only 18.06 and 55.56% of households access all weather road and market within five km distance, respectively, making the study communities isolated from market centers. In agreement with this finding, studies state that underdeveloped infrastructure is a driving cause for insufficient access to public services, minimal market integration and little returns on investments (30). Hence, geographically isolated communities who live distant from the main road and local market experience minimal access to inputs, market exchange, information as well as livelihood diversification opportunities [33, 27]. Likewise, Alinovi et al. (32) argue that access to basic infrastructure is determinant in promoting households’ resilience to shocks by enhancing their access to assets. Access to credit services was also minimal where only 59.38% of households access credit facilities in their proximity. Studies state that insufficient physical structures significantly limit access to basic services like health and credit facilities, contributing socioeconomic marginalization (33). In effect, lack of access to cash needs during crises is a major factor limiting households’ resilience to climate change-induced shocks (26).

### 3.2 Households’ resilience as measured by Climate Resilience Index and resilience capacities

The livelihood resilience analysis through the three-capacities and Climate Resilience Index showed relatively comparable results. Accordingly, the highland is better off in sociodemographic profile, water and health; the midland is better off in exposure to natural disaster and livelihood strategies and the lowland is better off in income and food access, asset, stability, social capitals and access to basic services (Annex 1; Table 3).

**Table 3.** Indexed major components, core-capacities and overall Livelihood Resilience Index of Dinki watershed socio-ecological system. (NDCV=Natural Disaster and Climate Variability; IFA=Income and Food Access; SDP=Sociodemographic Profile; LVS=Livelihood Diversity and ABS=Access to Basic Services).

|  |  | Agro-ecology |  |  |  |  |  |
| --- | --- | --- | --- | --- | --- | --- | --- |
|  |  | Highland |  | Midland |  | Lowland |  |
| Resilience capacities | Major components | Component value | Resilience score | Component value | Resilience score | Component value | Resilience score |
| Absorptive | NDCV | 0.472 |  | 0.657 |  | 0.503 |  |
|  | Stability | 0.45 | **0.448** | 0.414 | **0.517** | 0.412 | **0.520** |
|  | Social capital | 0.404 |  | 0.419 |  | 0.693 |  |
| Adaptive capacity | IFA | 0.412 |  | 0.491 |  | 0.516 |  |
|  | Health | 0.46 | **0.436** | 0.416 | **0.495** | 0.399 | **0.417** |
|  | Water | 0.544 |  | 0.465 |  | 0.361 |  |
|  | SDP | 0.569 |  | 0.455 |  | 0.459 |  |
|  | Assets | 0.288 |  | 0.31 |  | 0.371 |  |
|  | LVS | 0.343 |  | 0.444 |  | 0.385 |  |
| Transformative capacity | Social capital | 0.505 |  | 0.499 |  | 0.542 |  |
|  | ABS | 0.35 | **0.389** | 0.327 | **0.37** | 0.355 | **0.402** |
|  |  |  |  |  |  | Overall LRI | **0.444** |

The livelihood resilience analysis through resilience capacities more clearly differentiated the agro-ecological zones in terms of their absorptive, adaptive and transformative capacities. In effect, the leading contributing factor to the resilience of Dinki watershed socio-ecological system to climate change-induced shocks was observed to be absorptive capacity with a mean index value of 0.495 followed by adaptive capacity with a mean index value of 0.449 (Fig.2a). In terms of agro-ecology, the midland was found to be relatively more resilient to climatic shocks with a mean index value of 0.461 (Fig. 2b).

**Figure 2.**
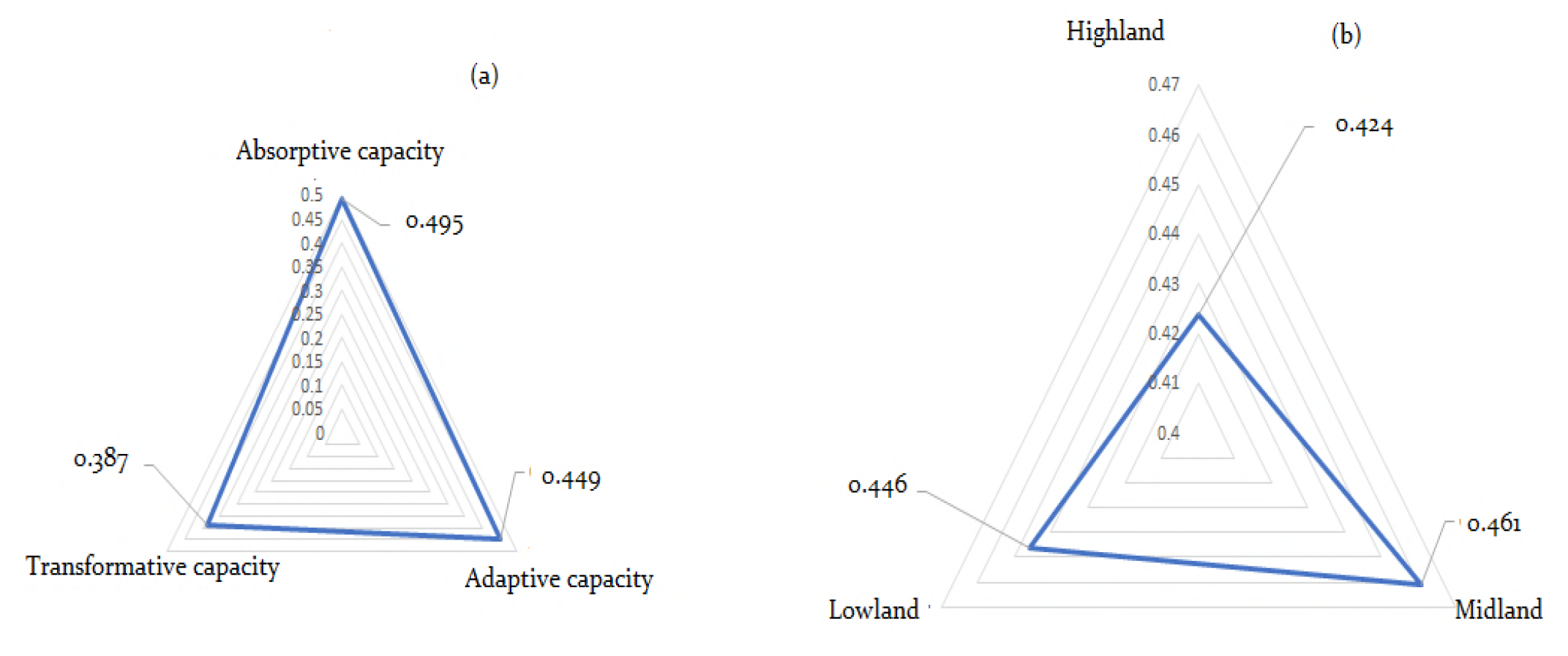
The resilience capacities (a) and resilience score of agro-ecological zones (b)

Relatively higher score of absorptive capacity in the lowland agro-ecology is evident by the fact that its exposure to recurrent climate change-induced shocks might have enabled residents to acquire more knowledge and get prepared for future likely shocks. Besides, large farm and livestock holding, social capital (CBOs, SSS, sharing of resources and technology) as well as coping strategies (in economic and management options) might have enabled lowland residents to better absorb shocks compared to the highland and midland agro-ecological zones.

In line with this study, Boka (34) disclose that households in Ethiopian lowland areas often have quick access to climate change information and early warning system contributing to their improved preparedness compared to other climatic zones. Other studies argue that large farm and livestock holding enable households to spread risks through income diversification and asset accumulation opportunities (27). Moreover, Frankenberger and his colleagues disclose that households’ ability to diversify financial capital, natural capital and social capital, among others, reduces their vulnerability, whilst enhancing their absorptive and adaptive capacities to properly respond to changing conditions (13).

On the other hand, the resilience score in terms of adaptive capacity was higher in the midland followed by the highland. It might be due to the fact that improved livelihood diversification practices (trade, irrigation, tree garden), technology utilization (improved seed and fertilizer) and improved access to credit might have enabled the midland and highland residents to better adapt climate change-induced shocks. Moreover, informal institutions like *idir* and *equib* are basic economic leverage contributing households to better adapt to shock impacts. In agreement with this finding, studies state that livelihood diversification, information exchange and economic leverage institutions contribute to enhance households’ adaptive capacity to shock impacts (13).

Although the mean resilience score in terms of transformative capacity (0.387) is lower to other resilience scores (Fig. 2a), the lowland showed the highest transformative capacity (0.402) than the other agro-ecological zones (Table 4). Relatively higher proportion of households who access market in their proximity coupled with higher social capital (transformative) scores through conflict management and vertical linkages in the lowland and highland might have contributed to higher transformative capacity in these agro-ecological zones. In this aspect, disputes over access to water, pasture and related land resources are repeatedly report as major sources of conflict in the study community. As a result, conflict management options through elders’ institutions might have contributed to build peace and security among the study communities.

In agreement with this finding, studies state that managing conflict ensures information exchange and market linkage with other communities leading to knowledge sharing. Besides, participation of community members in decision options facilitates information dissemination, access to basic assets during crises and enhance transformative capacity through institutional reforms (13). Furthermore, conflict management through customary laws are recognized as plausible options to sustain social capital among Africans (35). These institutions are participatory, easily accessible and sustainable in keeping peace and thereby resilience (13).

Furthermore, households’ resilience capacity was graphically presented in four quadrant charts following the Andersen and Cardona (36) and Weldegebriel and Amphune (25). The graph was established based on households’ income per capita and mean LRI values drawn on x and y-axes, respectively. Accordingly, based on the mean LRI value (0.44), households falling above the mean were poor but resilient, resilient and extremely resilient. Whereas rich but not resilient, vulnerable and extremely vulnerable households were presented below the mean. Likewise, based on the mean monthly income (18.66 per month or 0.622 USD per day), households falling to the right of the mean include rich but not resilient, resilient and extremely resilient. Whereas households who were poor but resilient, vulnerable and extremely vulnerable were presented in the left of the mean (Fig.3).

**Figure 3.**
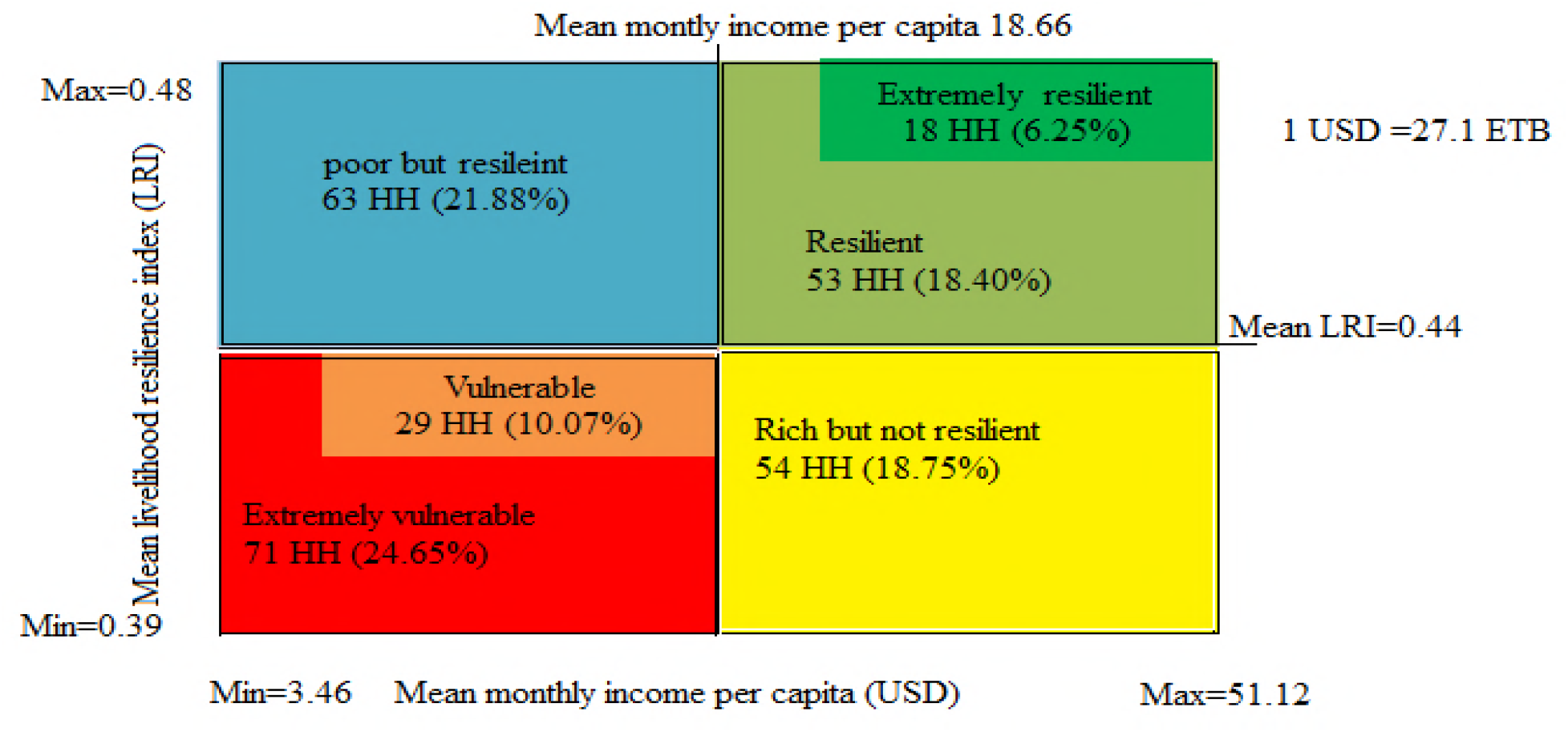
Resilience typologies by household monthly income

The average daily income value is far below the poverty line of sub-Saharan Africa, indicating the poverty level of the study communities. Moreover, even with this minimal cutoff, more than half of the households (56.59%) were vulnerable for poverty (Fig. 3). Factors, such as small asset holdings coupled with underdeveloped infrastructure might have limited their adaptive capacities signified by poor diversification practices, while amplifying their vulnerability.

In this study, considerable proportion (32.29%) of households own less than one hectare of land, nearly half (47.22%) of the study communities have less than two livestock unit and the overall infrastructure is underdeveloped (Annex 1). Moreover, the majority of households (94.79%) experience a single income-dominated livelihood options, making them vulnerable to climate change-induced shocks. In agreement with this finding, studies disclose that land and livestock are two of the most known financial assets in farming communities of Ethiopia critically determining their wealth status (27). Thus, poor households are often with small land size, few livestock unit and minimal livelihood options as well as are with minimal access to key household assets (natural, physical, human and social capitals) to diversify livelihoods and to empower their adaptive capacity (37). However, poor people are not necessarily vulnerable if have access to communication, infrastructure and support systems (38).

## 4. Conclusion

In this study, the Climate Resilience Index (CRI) and the resilience capacities (3Ds) frame were tested to measure households’ resilience to climate change-induced shocks. The methods presented detail description of factors contributing to households’ resilience to shock impacts. Likewise, access to and use of livelihood resources, such as farmlands, livestock, livelihood diversification, infrastructure, as well as social capital and ecological stability are identified to influence households’ resilience to climate change-induced shocks. However, it might be due to their exposure to recurrent shocks coupled with constrained adaptive capacities like limited diversification practices, poor access to infrastructure, underdeveloped social capital, among others, the mean resilience score of the study communities is minimal. Similarly, although improved absorptive capacity through early warning system, social protection, climate change information, etc. contributes to prepare, anticipate and cope with shock impacts, it is equally important to strengthen both the adaptive (adjustment strategies) and transformative (system-level change) capacities to ensure long-term resilience in the study communities.

## Acknowledgements

The authors would like to express their heart-felt gratitude to enumerators, study participants, and all who contributed to this study. All authors and sources of materials used in this paper are highly acknowledged.

---

**Annex 1.**
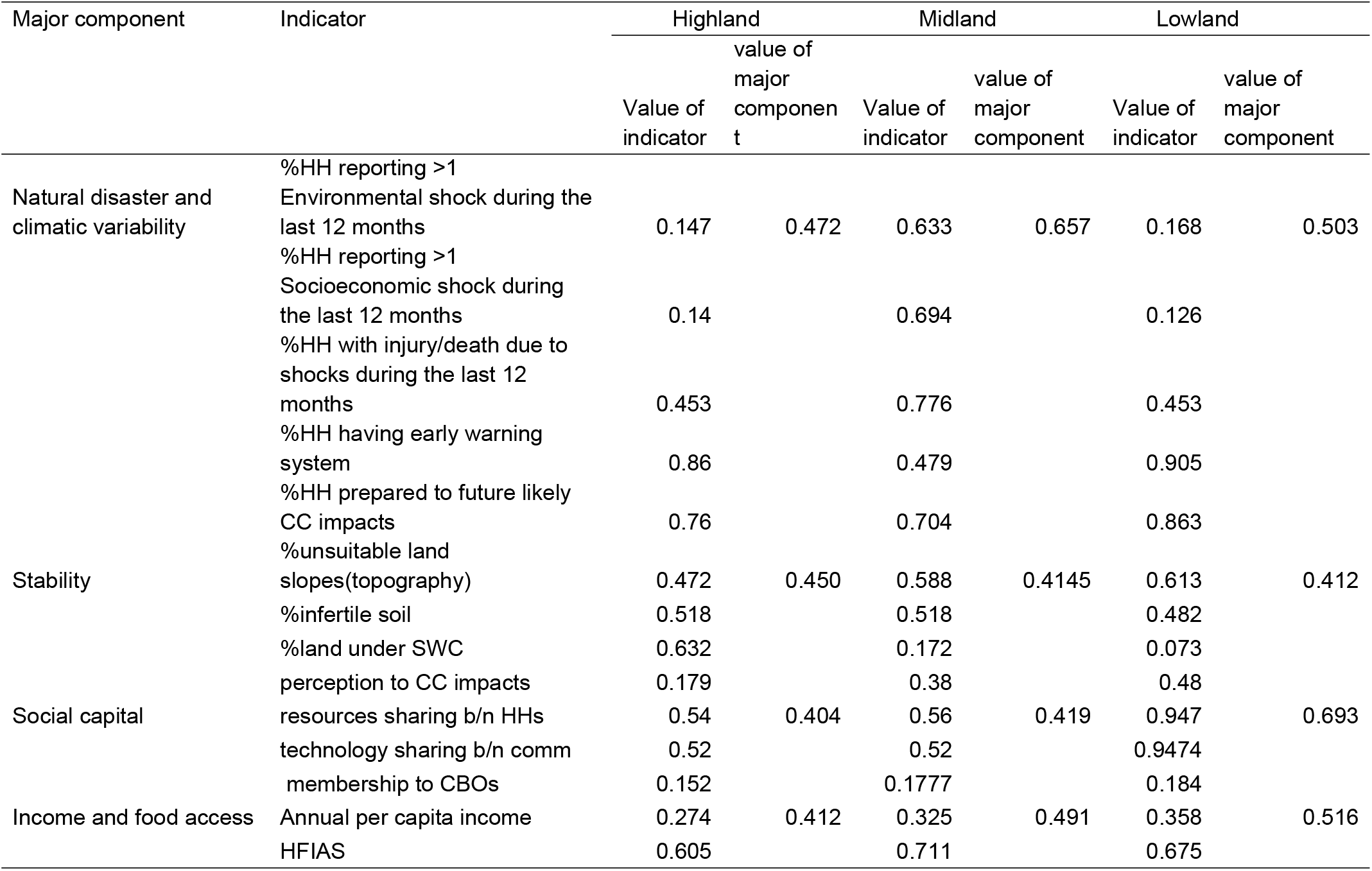
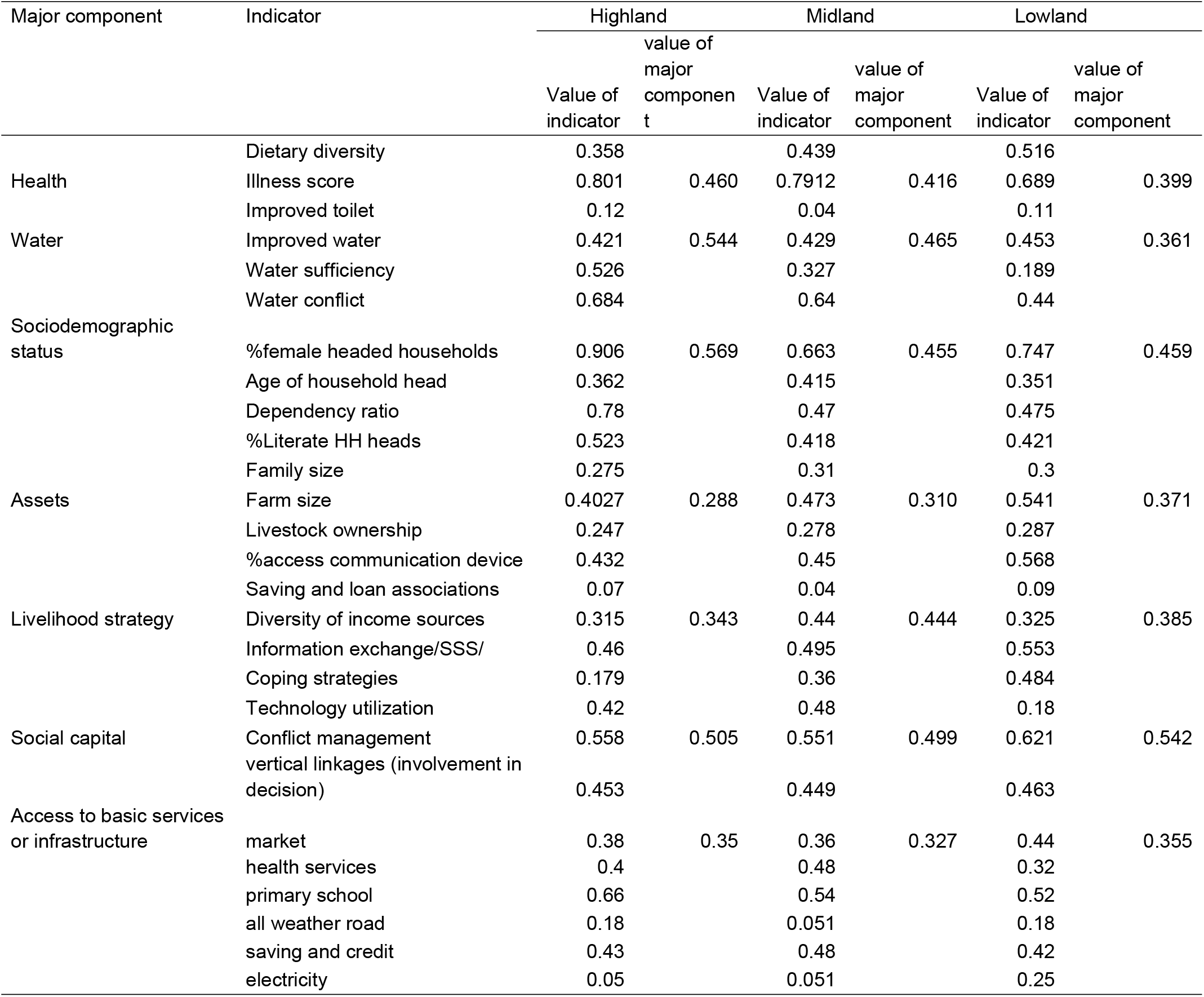
Indexed major components, sub-components and overall CRI of Dinki watershed socio-ecological system

